# Quantification of Early Gait Development: Expanding the Application of Catwalk Technology to an Infant Rhesus Macaque Model

**DOI:** 10.1101/2022.11.14.516450

**Authors:** Sabrina A. Kabakov, Emma Crary, Viktorie Menna, Elaina R. Razo, Jens C. Eickhoff, Natalie R. Dulaney, John R. Drew, Kathryn M. Bach, Aubreonna M. Poole, Madison Stumpf, Ann M. Mitzey, Kerri B. Malicki, Michele L. Schotzko, Kristen A. Pickett, Nancy J. Schultz-Darken, Marina E. Emborg, David H. O’Connor, Thaddeus G. Golos, Emma L. Mohr, Karla K. Ausderau

**Affiliations:** Department of Kinesiology, Occupational Therapy Program, University of Wisconsin– Madison, Madison, WI 53706, USA; Department of Pediatrics, School of Medicine and Public Health, University of Wisconsin– Madison, Madison, WI 53792; Department of Biostatistics and Medical Informatics, University of Wisconsin–Madison, Madison, WI 53792, USA; Department of Comparative Biosciences, University of Wisconsin–Madison, Madison, WI 53706, USA; Wisconsin National Primate Research Center, University of Wisconsin - Madison, Madison, WI, 53715, USA; Department of Obstetrics and Gynecology, University of Wisconsin–Madison, Madison, WI 53705, USA; Department of Medical Physics, University of Wisconsin – Madison, Madison, WI, 53705, USA; Department of Pathology and Laboratory Medicine, University of Wisconsin–Madison, Madison, WI 53705, USA; Waisman Center, University of Wisconsin–Madison, Madison, WI 53706, USA

**Author notes:** Corresponding author at Waisman Center, Department of Kinesiology, Occupational Therapy Program, University of Wisconsin–Madison, Madison, WI 53706, USA (K.K. Ausderau).

**Keywords:** Gait, Motor development, Rhesus macaque, Catwalk, Infant, Walking pattern

## Abstract

**Background:** Understanding gait development is essential for identifying motor impairments in neurodevelopmental disorders. Defining typical gait development in a rhesus macaque model is critical prior to characterizing abnormal gait. The goal of this study was to 1) explore the feasibility of using the Noldus Catwalk to assess gait in infant rhesus macaques and 2) provide preliminary normative data of gait development during the first month of life.

**New method:** The Noldus Catwalk was used to assess gait speed, dynamic and static paw measurements, and interlimb coordination in twelve infant rhesus macaques at 14, 21, and 28 days of age. All macaque runs were labeled as a diagonal or non-diagonal walking pattern.

**Results:** Infant rhesus macaques primarily used a diagonal (mature) walking pattern as early as 14 days of life. Ten infant rhesus macaques (83.3%) were able to successfully walk across the Noldus Catwalk at 28 days of life. Limited differences in gait parameters were observed between timepoints because of the variability within the group at 14, 21, and 28 days.

Comparison with existing methods: No prior gait analysis system has been used to provide objective quantification of gait parameters for infant macaques.

**Conclusions:** The Catwalk system can be utilized to quantify gait in infant rhesus macaques less than 28 days old. Future applications to infant rhesus macaques could provide a better understanding of gait development and early differences within various neurodevelopmental disorders.

**Highlights:**

- Infant rhesus macaque gait parameters can successfully be captured by the Catwalk
- At 14 days of life, macaques were consistently using a diagonal walking pattern
- Limited developmental change occurs in gait parameters over the first month of life
- Infant macaque gait parameters had high within group variability at each timepoint

**Graphical abstract:** 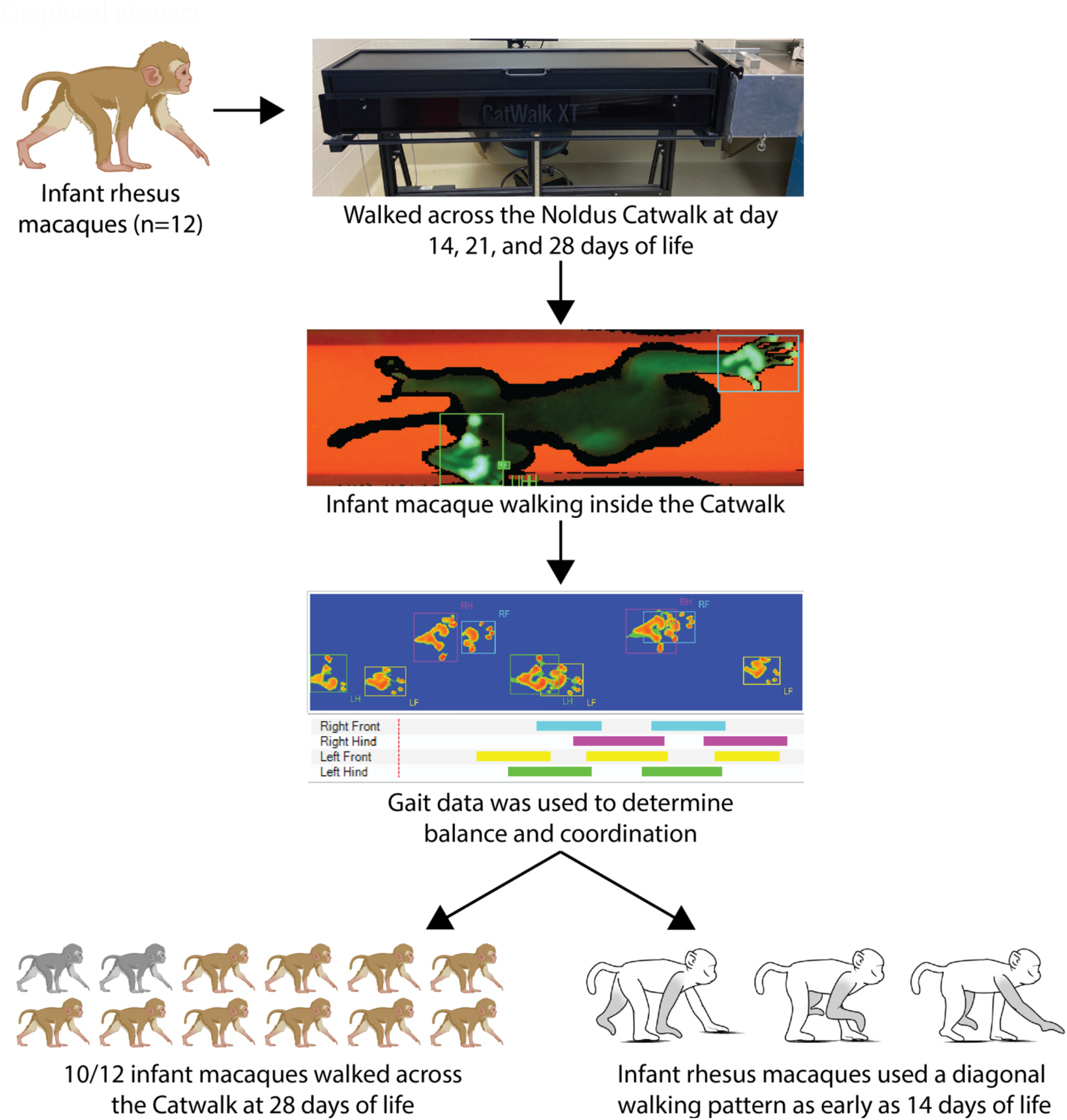

Note: Brown and grey-colored macaques were created in biorender

## 1. Introduction

Understanding gait development is essential for better identification and treatment of early motor delays in children. Infant gait emerges with a wide base of support and shorter steps to compensate for instability (Ivanenko et al., 2007). As children transition to a more mature gait pattern with less variability, their balance and control in the gait cycle increases as evidenced by decreased base of support and increased step length, speed, and time in swing phase (Adolph et al., 2003; Voss et al., 2020). Gait has been used as one measure that can differentiate neurodevelopmental disorders from typical development based on these gait parameters (Lum et al., 2021; Simmons et al., 2020) Motor deficits have been identified in children with autism spectrum disorder (Gawali et al., 2017; Green et al., 2009; Provost et al., 2007), attention-deficit/hyperactivity disorder (Lee et al., 2021), cerebral palsy (Gawali et al., 2017), developmental coordination disorder (American Psychiatric Association, 2013), and in general, intellectual and developmental disabilities (Bertapelli et al., 2020; Gawali et al., 2017). In addition, motor development can be influenced by insults such as prenatal Zika virus infection (Faiçal et al., 2019; Nielsen-Saines et al., 2019; Peçanha et al., 2020) or conditions causing preterm birth (Chung et al., 2020). Differences in motor development can be mild, including subtle differences in gait characteristics (Deconinck et al., 2006; Manicolo et al., 2019) and as severe as not achieving major motor milestones such as independent walking (Garfinkle et al., 2020; Melo et al., 2019). A nonhuman primate model could create a translational model of early motor development to identify motor differences within neurodevelopmental disorders to generate predictive measures for early motor deficits.

Rhesus macaques (*Macaca mulatta*) are the ideal model to study motor development as they more closely resemble human fetal brain and infant development compared to other animal models (Bauman & Schumann, 2018; Mohr, 2018). They also develop 3-4 times faster than humans allowing for longitudinal studies to be conducted in a shorter period (Dudley et al., 2019; Malkova et al., 2006). Studying rhesus macaque’s gait has been an effective method within spinal cord injury (Capogrosso et al., 2016), Parkinson’s disorder (Potts et al., 2014), and Huntington’s disease (Chan et al., 2015). By extending the rhesus macaque model into neurodevelopmental research, the subtle differences in gait development can be pinpointed and used as an important marker of motor impairments in humans.

Observational studies have been conducted describing typical infant rhesus macaques’ walking patterns. Most quadrupedal animals adopt either a lateral or diagonal gait pattern. Lateral gait is defined as when ipsilateral limbs move through the air and land at relatively the same time, while diagonal occurs when the contralateral limbs move together (Dunbar, 1989; Shapiro et al., 2014). Infant rhesus macaques begin ambulating 3-14 days after birth using a lateral gait pattern, which is classified as an immature walking pattern (Castell & Sackett, 1973; Hildebrand, 1967).

Between days 18 to 42, balance and coordination continue to improve, which contributes to a more mature gait and transition into an efficient diagonal walking pattern (Dunbar, 1989; Hildebrand, 1967). These observational studies included small sample sizes and relied on observational measures, lacking quantitative measurements needed to understand the specificity of gait patterns in rhesus macaques. It is currently unknown how specific gait parameters (step length, base of support, speed, and many others) change during the first month of life in infant rhesus macaques. There is a need for an objective gait analysis quantification system to measure gait development and variability in a rhesus macaque model.

Currently there are no computerized systems for categorizing gait development in infant rhesus macaques. Extending the application of the Noldus Catwalk XT 10.6 (Noldus Information Technology, Wageningen, the Netherlands) to rhesus macaques would create an objective measurement of gait. The Catwalk was originally designed to measure gait in rodents and has been successfully applied to nonhuman primate models of adult common marmosets (Pickett et al., 2020) and gray mouse lemurs (Le Corre et al., 2018). We hypothesized that the Catwalk system may be a viable tool to examine infant rhesus macaque gait development with the intent of establishing a procedure and normative measures for use in future translational studies. The purpose of this study was to examine the feasibility of collecting gait data in infant rhesus macaques and to quantify preliminary normative gait development over the first month of life.

## 2. Methods

### 2.1 Subjects and animal care

The twelve female infant Indian-origin rhesus macaques (*Macaca mulatta*) were part of the control arm of a larger prenatal Zika virus infection study at the Wisconsin National Primate Research Center (WNPRC). All animals were housed in appropriate enclosures of the Specific Pathogen Free (SPF) colony and were free of Macacine herpesvirus 1 (Herpes B), simian retrovirus type D (SRV), simian T-lymphotropic virus type 1 (STLV), and simian immunodeficiency virus (SIV). Dams were provided a dry diet that contained adequate caloric energy, carbohydrate, fat, protein, fiber, mineral, and vitamin content. The diet was supplemented with enrichments and additional food (fruit, vegetables, and treats) to encourage foraging. All rhesus macaques involved in this project were cared for by the staff at WNPRC and observed at least two times throughout the day by animal care staff for their appetite level, stool quality, activity level, and physical condition to discern any signs of pain, distress, and illness. Any animals with abnormal observations were evaluated by veterinarians to provide necessary care. Regulations and guidelines were followed according to the Animal Welfare Act and the Guide for the Care and Use of Laboratory animals, the principles described in the National Research Council’s Guide for the Care and Use of Laboratory Animals (National Research Council (US) Committee for the Update of the Guide for the Care and Use of Laboratory Animals, 2011), and the recommendations of the Weatherall report (Weatherall, 2006). The University of Wisconsin-Madison, College of Letters and Science and Vice Chancellor for Research and Graduate Education Centers Institutional Animal Care and Use Committee approved the nonhuman primate research covered under protocols G006108 and G005401.

These infants were born to dams that were inoculated with phosphate buffered saline at 30-45 days of gestation (GD). Prior to and after inoculation, the dams were monitored daily for any clinical signs of illness (e.g., diarrhea, inappetence, inactivity, fever and/or atypical behaviors). As part of the larger study, pregnant dams were sedated every 1-2 weeks for a prenatal ultrasound to monitor fetal health and had blood draws weekly throughout pregnancy. All infants were scheduled to be delivered by cesarean section at about 160 GD. Infants were delivered 6 days earlier than the average natural birth at WNPRC of 166 GD to ensure collection of the placentas for the larger research project. One animal was naturally born (044-508) at 159 GD prior to their scheduled delivery. All infants were dried off, stimulated, and Apgar scores were assigned as previously described (Ausderau et al., 2021) (Table 1 and Supplemental Table 1).

**Table 1.** Infant characteristics

|  | Mean (95% Confidence Interval) |
| --- | --- |
| Gestational age at saline inoculation (days) | 39.8 (34.1-45.5) |
| Delivery method by Cesarean section (%) | 91.7 (73.3-100) |
| Female sex (%) | 100(100-100) |
| Gestational age at birth (days) | 159.8 (158.9-160.6) |
| Weight (kg) |  |
| Birth weight | 0.51 (0.47-0.54) |
| 14 days | 0.50 (0.45-0.55) |
| 21 days | 0.53 (0.47-0.59) |
| 28 days | 0.56 (0.49-0.63) |
| Apgar scores |  |
| 1 minute | 5.9 (4.9-6.9) |
| 5 minutes | 7.1 (6.2-8.0) |
| 10 minutes | 8.6 (7.8-9.5) |
| Housing status with dam or surrogate at 1 month (%) | 75 (46.2-100) |
| Age at gait testing |  |
| Age at timepoint 1 (days) | 14.5 (13.8-15.2) |
| Age at timepoint 2 (days) | 21.2 (20.6-21.8) |
| Age at timepoint 3 (days) | 28.4 (27.7-29.2) |

Some infants were rejected by their mothers and alternative housing was attempted with a surrogate dam. Three animals remained in the nursery for the entire study. Infants who remained in the nursery were fed 5% dextrose then formula. Animals immediately housed with their biological or surrogate mothers were not provided supplemental formula. Infants were only removed from their cage for attempted pairings with a surrogate or for neurodevelopmental and gait assessments during the first month of life.

### 2.2 Data Collection

Infants were removed from the nursery or separated from their mothers for neurodevelopmental testing with the Schneider Neonatal Assessment for Primates (SNAP) and Catwalk assessment. The SNAP test was first, taking approximately 30 mins to test the infant’s motor, orientation, sensory responsiveness, and state control (Ausderau et al., 2021). The infant’s performance on the SNAP motor test was used to provide additional insight into the infant’s ability to complete the gait assessment using the Catwalk.

The Catwalk XT version 10.6 was modified for infant rhesus macaques in accordance with the Pickett et al. (2020) changes for common marmosets. The original goal box at the end of the Catwalk was removed and sliding tracks were added to allow a nest box to slip on and off the end. The nest box was used to transport the rhesus macaque to the Catwalk and was placed on the end for the infant macaque to have a closed cage to walk into. The walls of the Catwalk were adjusted to 10,8cm to limit lateral movements and reduce variability in walking. The camera was placed at 66.5cm from the floorplate to optimize visibility of the walkway glass. The detection settings were consistent between trials with the camera gain 20.0dB, green intensity threshold 0.14, red ceiling light 17.7V, and green walking light 16.0V. The testing room that contained the Catwalk had limited stimulation and was quiet to keep the infant calm throughout the assessment. In addition, the lights in the room were off to ensure optimal brightness of the LED lights on the walkway. All testing occurred at a similar timeframe to adjust for daily routines to account for circadian rhythm.

Research team members directly handled and placed the infant into the opening of the CatWalk where the infant ambulated through a 130 cm long walkway (Figure 1). As the animal moved across and applied pressure on the walkway, the green LEDs were refracted on the red background and individual footfalls were captured with a high-speed digital camera. At the end the walkway, the infant entered the nesting box or was picked up by a research team member.

**Figure 1.**
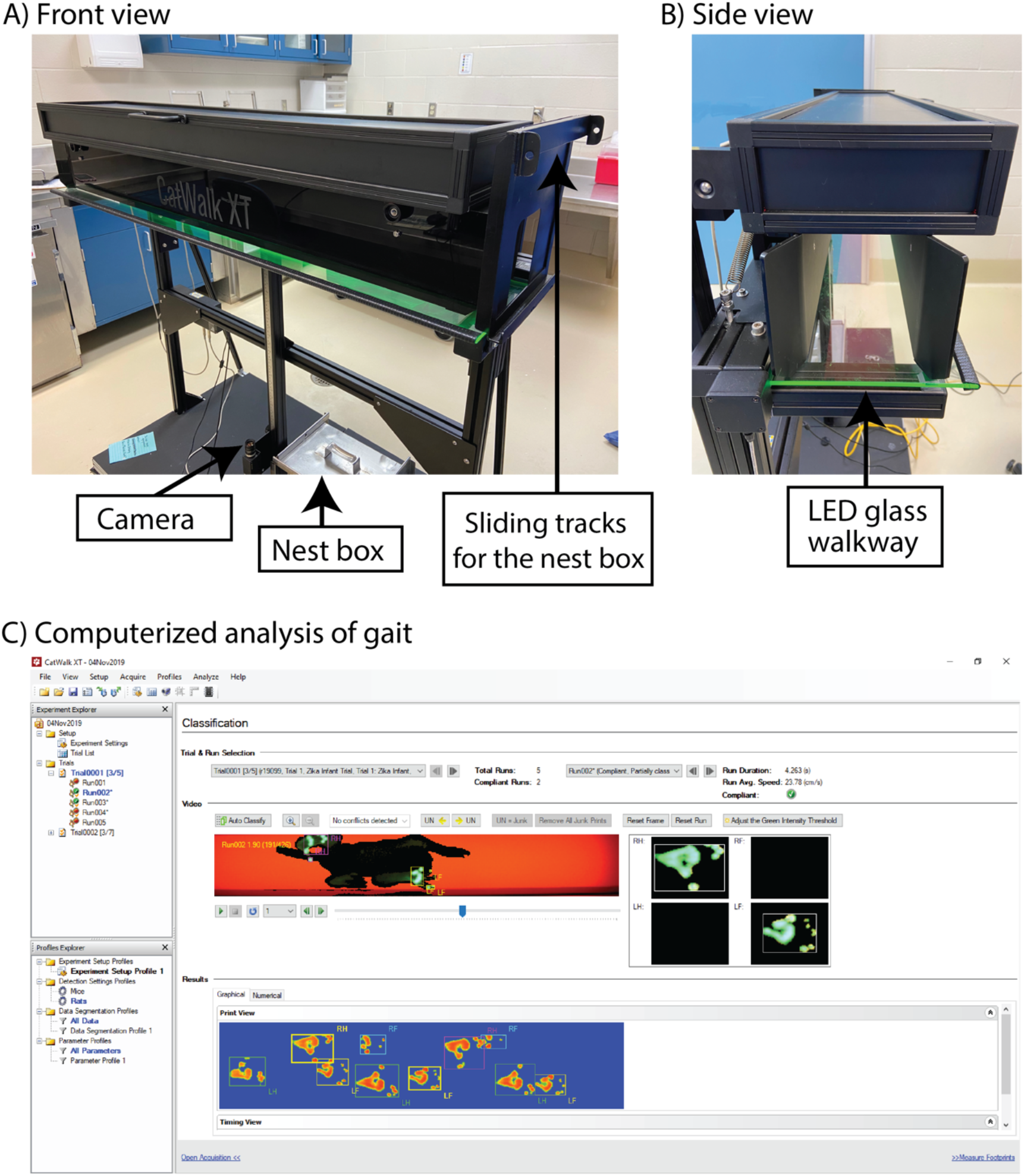
Catwalk apparatus A) Front view of Catwalk apparatus with the LED lights on. B) Side view of Catwalk where the infant macaque enters the walkway. C) Output on the computer showing the infant in the Catwalk with limbs categorized.

The criteria for a usable run were established based on Pickett et al. (2020) where each infant must have at least two consecutive footfalls per limb on the walkway without stopping or jumping/leaping (Video 1). The infant was tested until at least three usable runs were collected, or the infant was unable or unwilling to complete the task. If the infant did not complete a usable run, the tester attempted to train the infant by putting them partially through the Catwalk system or placing their blanket in the system as a reinforcer. Data were collected on days 14, 21, and 28 ± 1 day of life. The infant’s birth was classified as day 1 of life.

**Video 1.**
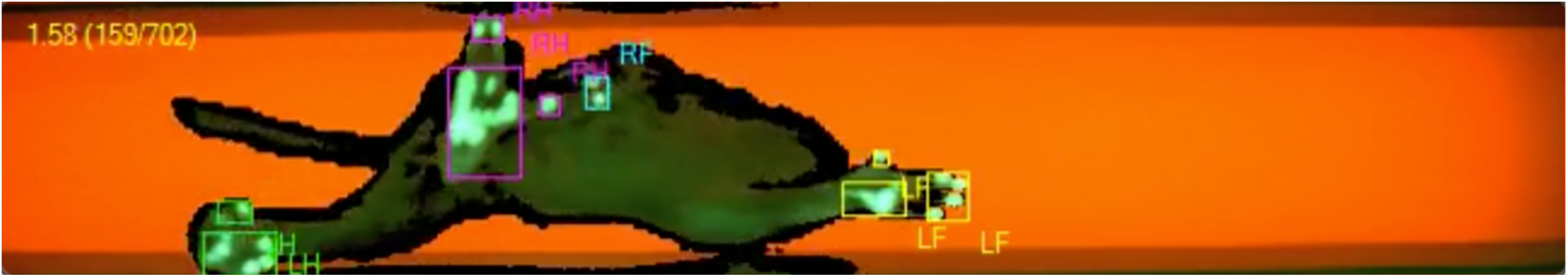
Example of a rhesus macaque walking through the Catwalk apparatus

## 2.3 Data Preparation

All runs used an adjusted green intensity threshold of 0.31 to reduce any extra noise when labeling the limbs. Recorded infant runs were reviewed by two independent coders to determine which ones were usable with all limbs being labeled as right front (RF), left front (LF), right hind (RH), and left hind (LH). Any errors or disagreements in coding were decided by a group consensus. Feasibility to complete the task was measured by the animal’s ability to successfully move across the Catwalk walkway without stopping.

Catwalk software calculated the trial statistics, an average (mean) of the three runs, for each time point for all variables. The Noldus technical support team was consulted to apply the Catwalk software most effectively to an infant rhesus macaque model and support selection of the appropriate analysis variables. We selected gait and paw variables used in the Catwalk analysis that were predicted to measure the development of balance and coordination. The variables selected were categorized into speed, dynamic paw measurements, static paw measurements, and inter-paw coordination. Selected variables are listed in Table 2, including definitions, limb output, and equations (Figure 2). The percentage of time standing on limbs was calculated using the raw data as compared to the Catwalk software due to insufficient number of infant rhesus macaque footfalls on the walkway. The calculation started when all four limbs were visible within the walkway and ended when at least one limb was no longer visible.

**Table 2.** Gait variables

| Gait variable (units) | Definition<sup>1</sup> | Limb Outputs | Equation |
| --- | --- | --- | --- |
| *Speed* |  |  |  |
| Speed (cm/s) | How fast the animal walked across the walkway | N/A | $\frac{Distance\ (cm)}{Time\ (s)}$ |
| Stand index (arbitrary units) | Reported as a negative value to reflect the speed at which the limb is lifted off the walkway starting from the point of max contact to 90% of contact being off the walkway | RF, RH, LF, LH | $\frac{a^{\#}}{max\ contact\ area} \times FR^{\#\#}$ |
| *Dynamic paw measurements* |  |  |  |
| Step cycle (s) | Time between two consecutive initial contacts of the same limb used as common denominator for dynamic paw measurements to remove the impact of time | RF, RH, LF, LH | N/A |
| Stand (s) | Total time one limb is in contact with the glass walkway | RF, RH, LF, LH | N/A |
| Duty cycle (%) | Percent of time the infant had their limb on the walkway, stand | RF, RH, LF, LH | $\frac{Stand\ (s)}{Step\ cycle\ (s)} \times 100$ |
| Swing (%) | Percent of time the infant had their limb in the air | RF, RH, LF, LH | $\frac{Swing\ (s)}{Step\ cycle\ (s)} \times 100$ |
| Single stance (%) | Percent of time the front or hind limb is standing on the walkway without the contralateral limb (ex. LF is in contact with the walkway without RF) | RF, RH, LF, LH | $\frac{2^*}{Step\ cycle(s)} \times 100$ |
| Dual stance (%) | Percent of time both the front or hind limbs are in contact with the walkway (ex. LF and RF are in contact with the walkway) | Front or hind | $\frac{(1+3)^*}{Step\ cycle\ (s)} \times 100$ |
| *Static paw measurements* |  |  |  |
| Stride length (cm) | Distance between center placements of the same paw. | RF, RH, LF, LH | Pythagoras' theorem |
| Base of support (cm) | Distance between the placement of the left paw to the right paw | Front or hind | N/A |
| Print position (cm) | Distance between where the ipsilateral hind paw is placed in relation to the front paw in one step cycle<br>+ print position= hind paw is placed behind the front paw<br>- print position= hind paw is in front of the front paw | Left or right | N/A |
| *Inter-paw coordination* |  |  |  |
| Phase dispersion cstat mean (%) | Temporal measurement between the initial contact of an anchor paw to the target paw touching the walkway<br>0% or 100% = anchor and target paw touch the walkway at the same time<br>50% = target paw lands halfway through the anchor limbs step cycle. | (anchor → target)<br>RF → LH, LF → RH,<br>LH → RH, LF → RF,<br>RF → RH, LF → LH | $\frac{(ICT-ICA)^{**}}{\text{Step cycle anchor (s)}} \times 100$ |
| Phase dispersion diagonal walking pattern | Categorization of phase dispersion cstat mean into a dichotomous outcome of diagonal or non-diagonal pattern based on the mean $\pm 1$ SD<br>Expected values (Figure 2C):<br>Contralateral limbs (RF → LH, LF → RH) = 0/100% $\pm 1$ SD<br>Girdle limbs (LH → RH, LF → RF) and ipsilateral limbs (RF → RH, LF → LH) = 50% $\pm 1$ SD | RF → LH, LF → RH,<br>LH → RH, LF → RF,<br>RF → RH, LF → LH | N/A |
| Standing on limb (%) | Percent of time the infant is standing on a combination of paws during the entire run | Contralateral (RF-LH or LF-RH), three or four limbs | $\frac{X^{***}}{\text{total \# steps}} \times 100$ |

<sup>1</sup>Definitions were created using the Noldus Catwalk manual and modified based on the application in this manuscript. $a^{\#}$ is derived from $y=ax+b$ describing the best fit line from max contact to 90%. $FR^{\#}$ is the frame rate.
\*= See Figure 2A for further explanation of numbers 1, 2, and 3 in the equation. \*\*= Initial contact time of the target limb- Initial contact time of the anchor limb. X\*\*\*= zero, one, diagonal, ipsilateral, girdle, three, or four.

**Figure 2.**
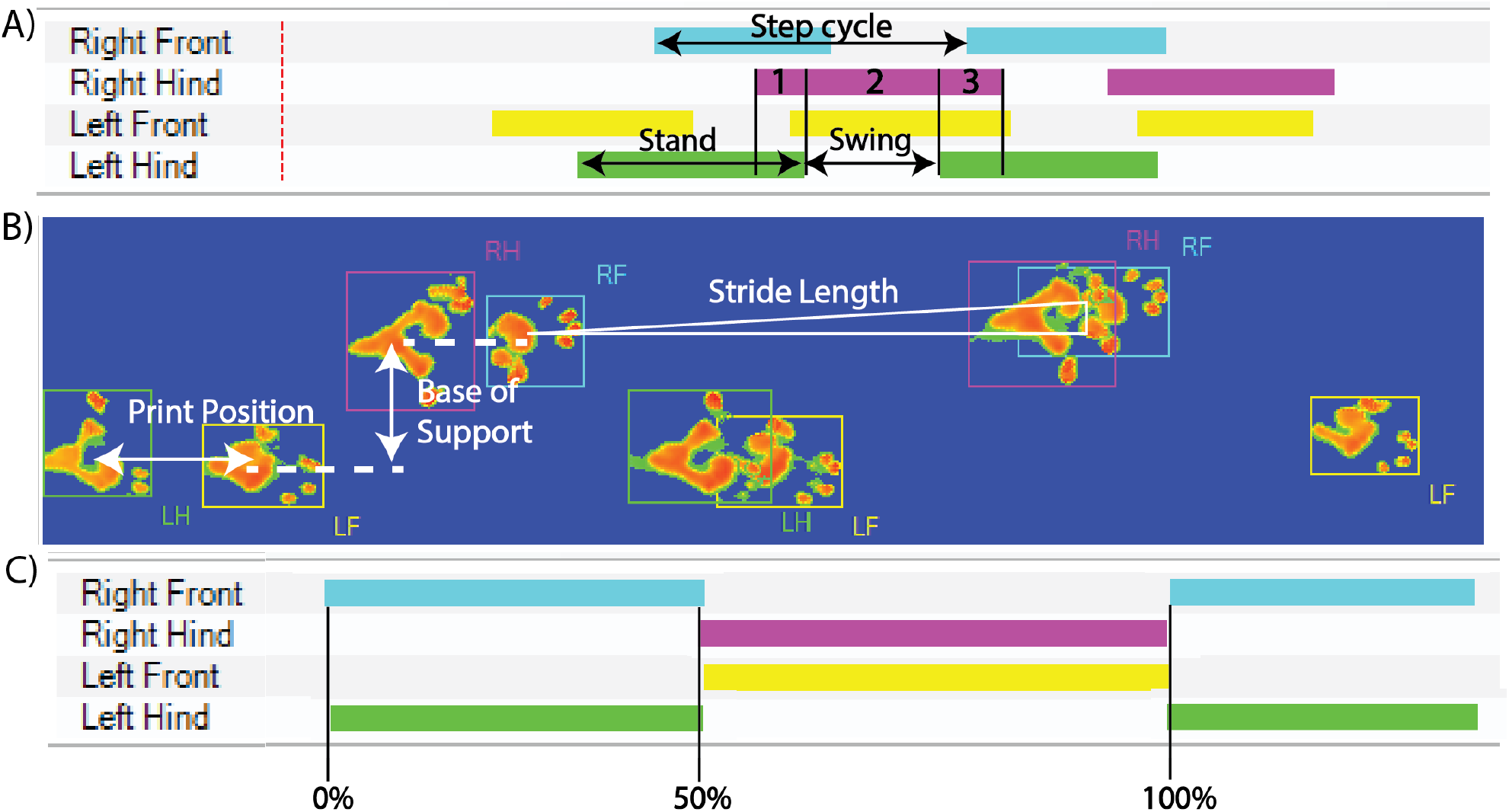
Visual representation of Catwalk variables. A) Represents the dynamic paw measurements showing stand (duty cycle), swing, single stance (2), and dual stance (1+3). Step cycle, time between two consecutive footfalls of the same limb, is used to create percent of time in the dynamic variables. B) Static paw measurements. Stride length was measured using the hypotenuse to calculate the distance between the center of two consecutive limbs. Print position and base of support are visually represented using the center of the print. C) Represents a diagonal phase dispersion pattern with the RF limb as the anchor limb and the RH, LF, and LH being the target limb. Diagonal phase dispersion will have RF and LH moving at the same time as represented by 0 and 100%, while the RH and LF will land about halfway into the step cycle of RF resulting in a cstat mean of 50%.

### 2.4 Categorizing walking pattern

To ensure the objective gait data reflected the walking pattern, all the infant rhesus macaque runs were reviewed and categorized as a diagonal or non-diagonal pattern. A diagonal walking pattern was defined as contralateral forelimbs and hindlimbs moving close together in time during the swing and stand phase (Shapiro et al., 2014) (Figure 3). A non-diagonal pattern was defined as any other combination of limbs including lateral when the ipsilateral forelimb and hind limb swing and land close together in time, when only one limb is moving at a time, and if a mixed combination of a diagonal and non–diagonal pattern was observed. All runs were observed by at least two independent coders to determine if the infant used a diagonal, mixed, or non-diagonal walking pattern. Disagreements in walking pattern classification were discussed until consensus was reached by the research team.

**Figure 3.**
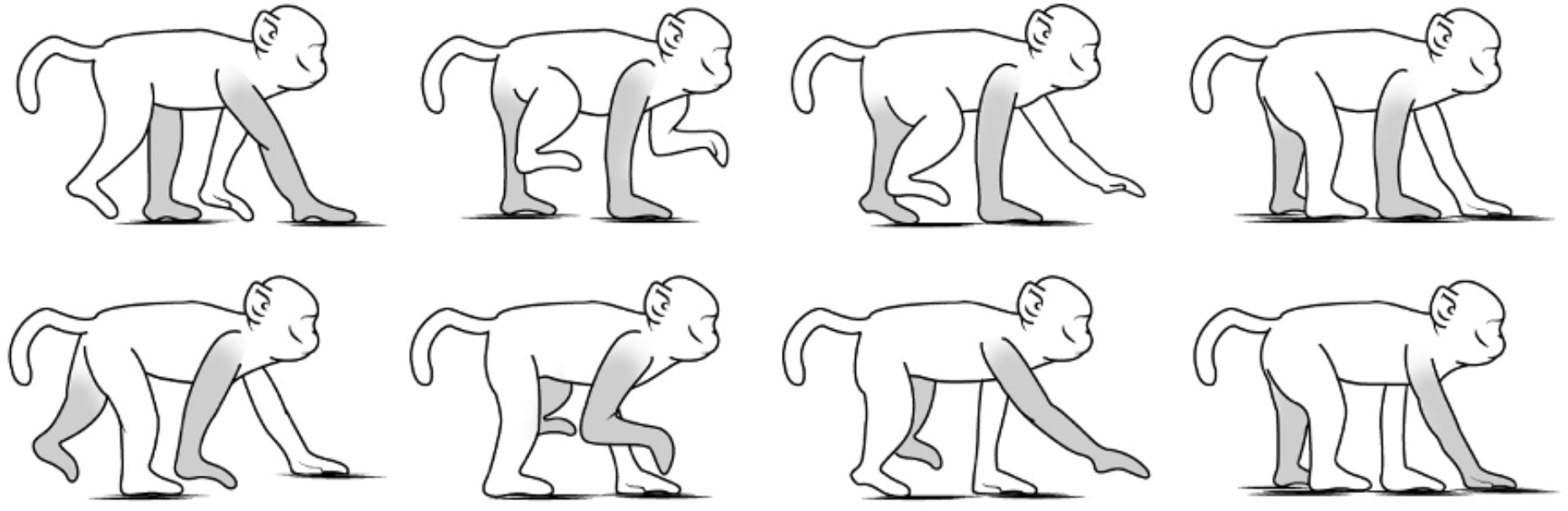
Visualization of an infant macaque using a diagonal walking pattern. The RF and LH limbs are shaded in gray to follow one full step cycle. The contralateral limbs are moving through the stand (limbs on the walkway) and swing (limbs are in the air) phase close together in time.

## 2.5 Data Analysis

Statistical analyses were conducted using SAS software (SAS Institute, Cary NC), version 9.4 and graphical figures were made using R statistical software (R Core Team, 2019) using the package ggplot2 (Wickham, 2016). Continuous variables were summarized using means and confidence intervals (CIs). Categorical characteristics were expressed as frequencies and percentages. Animal Catwalk success rates at each assessment time point were summarized descriptively in tabular format and compared between assessment time points using Fisher’s exact test. Test-retest reliability of Catwalk parameters was evaluated by examining the intra-class correlation coefficient (ICCs) at each timepoint. Gait outcome parameters were assessed at days 14, 21 and 28, and analyzed for each limb using linear mixed effects model with animal specific random effects and an autoregressive correlation structure to account for correlations between repeated measurements within each animal. The analyses were adjusted for birthweight and days before placement with a dam as nursery-reared animals during the first month engage in more locomotion than mother-reared infants (Champoux et al., 1991; Harlow et al., 1963). Model adjusted means and the corresponding 95% CIs were reported for each assessment time point. Walking pattern and phase dispersion outcomes were categorized as diagonal vs. non-diagonal pattern and analyzed using a generalized linear mixed effects model with animal specific random effects and a logit link function. All reported p-values are two-sided and P<0.05 was used to define statistical significance.

## 3 Results

### 3.1 Rhesus Macaque Catwalk Feasibility and Reliability

Twelve animals were assessed on their ability to complete at least one run across the Catwalk with usable data (at least two consecutive footfalls per limb on the walkway without stopping or jumping/leaping). The infant’s ability to perform the task at day 14 was the lowest (66.7%), increased at day 21 (75%), and by day 28 most infants (83.3%) had at least one usable run. There were no significant differences between the proportion of infants able to complete at least one run at these different time points (Supplemental Table 2). Infants unable to complete the task at day 21 and day 28 were reported to be resistant to going into the Catwalk apparatus or distracted by the environment while infants at day 14 appeared to have difficulty with the motor skills. We explored the test-retest reliability of continuous variables between runs for each infant within a timepoint with most variables demonstrating poor to moderate reliability (Supplemental Table 3).

### 3.2 Preliminary normative data

The second goal of this study was to examine differences in gait parameters at 14, 21, and 28 days of life to provide preliminary normative data for infant rhesus macaques. We calculated the means and 95% CI (Table 3) for each day after adjusting for the infant’s number of days in nursery before placement with a dam and birthweight.

**Table 3.**
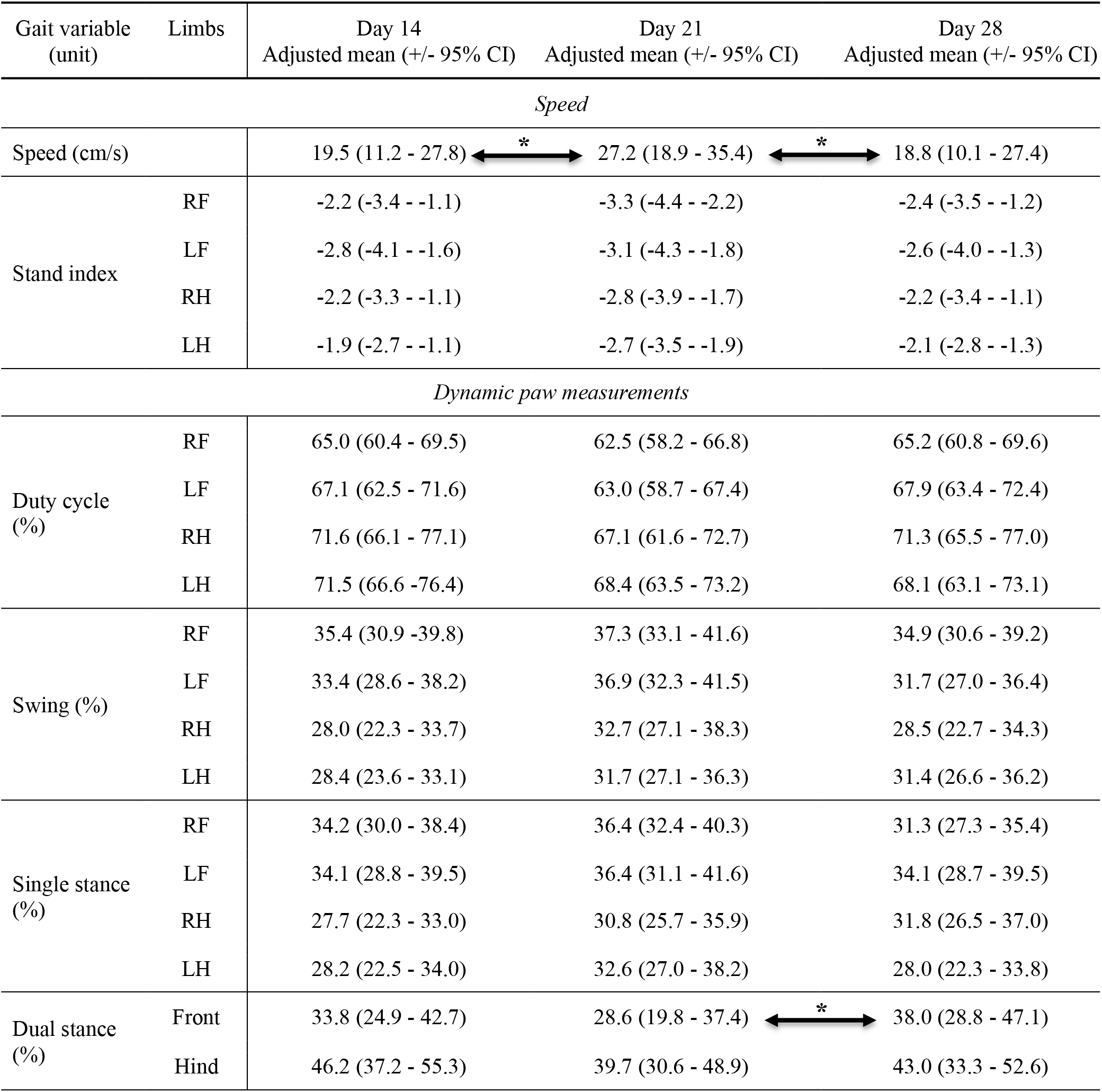
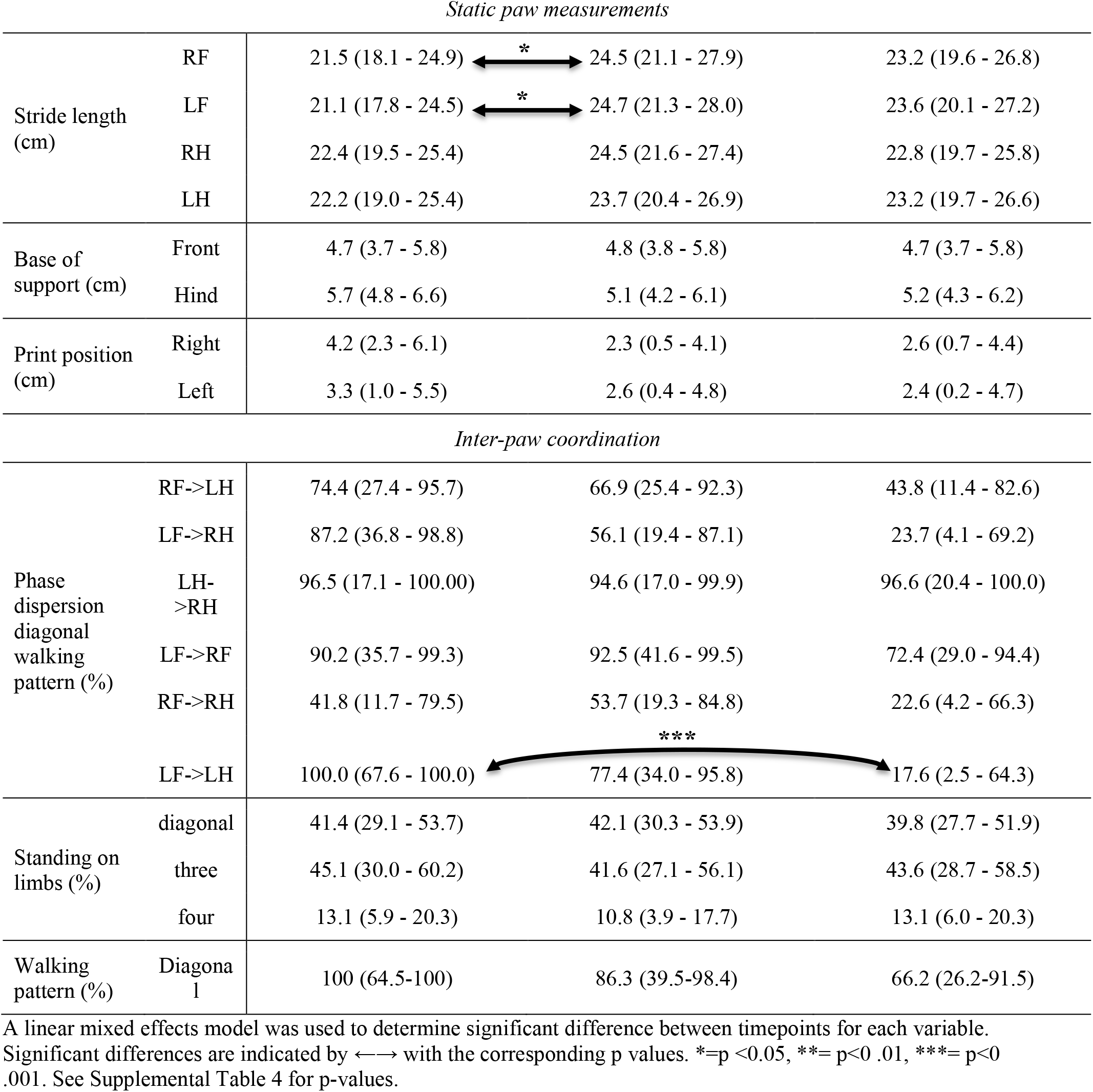
Catwalk variable data

Speed increased significantly from day 14 to day 21 (p=0.049) and then decreased from day 21 to day 28 (p=0.032), there was no difference between day 14 and day 28 (Figure 4A). The percent of time that the infant maintained both front paws on the walkway, i.e. dual stance, increased significantly from day 21 to day 28 (p=0.035) with no noticeable change from day 14 to day 21 (Figure 4B). Front stride length increased significantly from day 14 to day 21 (LF p=0.019; RF p=0.049), and then stayed steady at day 28 (Figure 4C, D) Dynamic paw measurements and static paw measurements did not differ significantly from day 14 to 28 of life (Supplemental Table 4). There were no significant differences in standing on limbs (%) with the infants spending about the same amount of time on diagonal limbs, three and four limbs between day 14 to day 28 of life. There was a widespread variability at each time point as represented by the large ranges of confidence intervals (Table 3).

**Figure 4.**
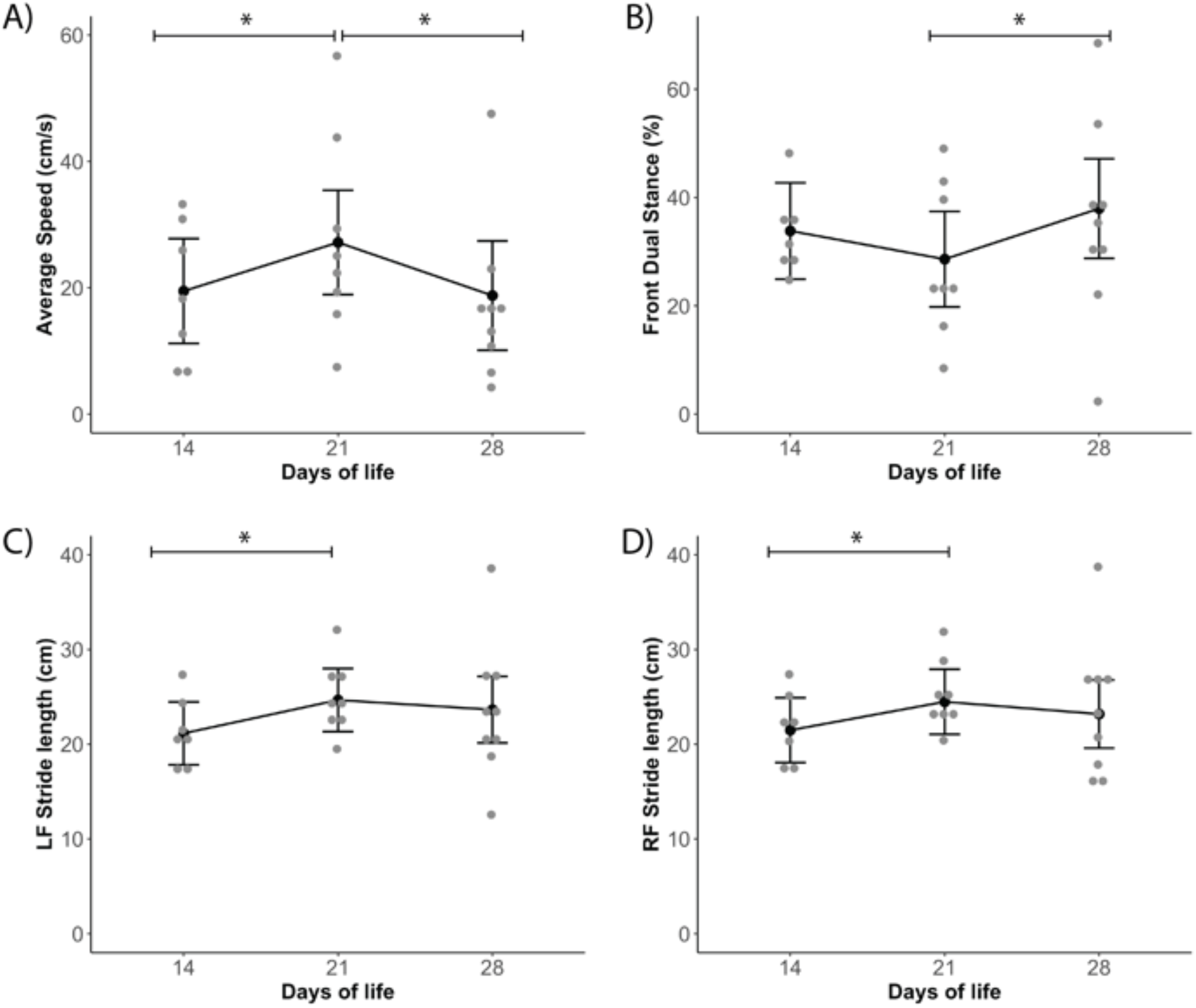
Catwalk parameters with significant changes over time. Adjusted means for days without a dam and birthweight ± 95% confidence interval is graphed for all variables. A line with star represents significant differences between timepoints. The gray dots represent individual data points. A) Speed, B) Front dual stance, C) LF Stride Length, and D) RF stride length. *=p <0.05, **= p<0 .01, ***= p<0 .001.

### Walking Pattern

A diagonal walking pattern, where the contralateral front and hind limbs move synchronously, is considered a developmentally mature walking pattern for rhesus macaques (Shapiro et al., 2014). We compared phase dispersion parameters, which represents the time between when the anchor and target limbs are placed on the walkway. In a mature diagonal walking pattern, we expected phase dispersion parameters to have contralateral limb pairs with a cstat mean of 0 or 100% and ipsilateral and girdle limb pairs to be at 50%. Phase dispersion was converted into a categorical variable of the anchor and target limb either using or not using the diagonal pattern based on their cstat mean landing within expected range (Figure 5). We found one significant difference with the LF→ LH pair having 100% of the limb pair timing falling within the diagonal phase dispersion at day 14 of life and only 17.6% at day 28 of life (p=0.0002). At day 14 of life, all the limb pairs, except for the RF→ RH pair, had their timing falling within the diagonal walking patten. These findings suggest infant rhesus macaques are using a diagonal walking pattern around day 14.

**Figure 5.**
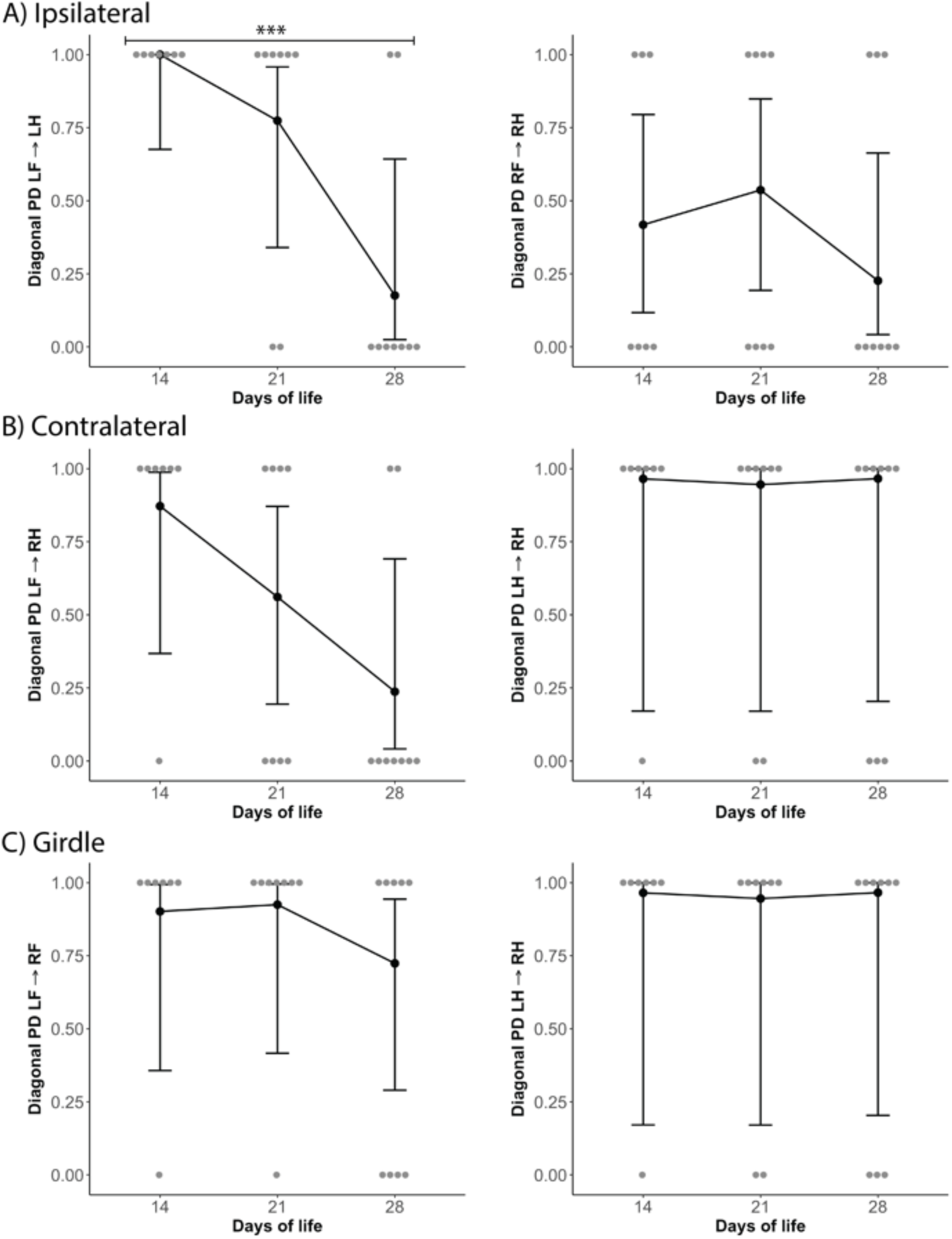
Phase dispersion (PD) diagonal walking pattern. Diagonal phase dispersion was a dichotomous variable of diagonal (1) or non-diagonal (0). The gray dots represent the individual animals, and the black dots are the adjusted mean ± 95% confidence interval. A) Ipsilateral limb pairs (LF→ LH, RF→ RH), B) contralateral limb pairs (LF→ RH, RF→ LH), and C) girdle limb pairs (LF→ RF, LH→ RH). *=p <0.05, **= p<0 .01, ***= p<0 .001.

We found no significant differences in the percent of animals using a diagonal walking pattern between day 14, 21, and 28 (Figure 6). Observers identified slight variation in timing in diagonal walking patterns for 50% of the infant’s Catwalk runs. The front limb typically landed slightly earlier than the hind limb but still used a diagonal pattern of the contralateral limbs moving together through stand and swing phase.

**Figure 6.**
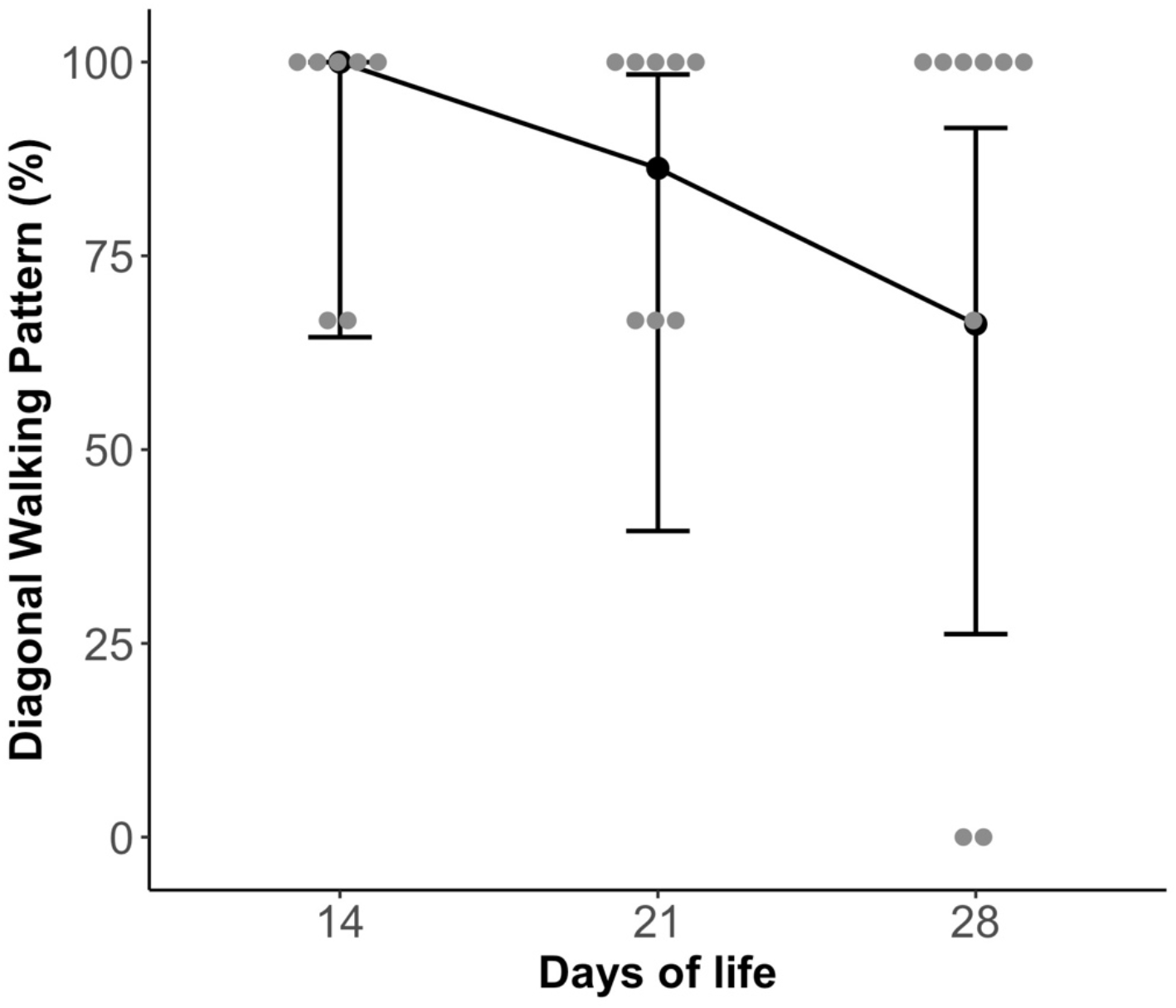
Percent of infant macaques using a diagonal walking pattern. There was a non-significant decrease in diagonal walking pattern usage by day 28 of life. Individual data is graphed in gray representing the percent of the usable runs that were categorized as a diagonal pattern. The adjusted mean based on days before placement with a dam and birth weight ± 95% confidence interval is graphed in black.

### 4. Discussion

The purpose of this study was to expand the application of the Noldus Catwalk XT into a rhesus macaque model and define preliminary normative gait development at 14, 21, and 28 days of life. This study established that it is feasible to use the Noldus Catwalk to collect gait data on infant rhesus macaques in the first month of life. The technology provided a quantitative classification of typical walking showing limited changes over time in overall motor developmental coordination and balance variables during the first month of life. However, 100% animals were identified to be using a mature diagonal walking pattern at 14 days of life, which is earlier than previously reported in the literature (Castell & Sackett, 1973; Dunbar, 1989; Hildebrand, 1967).

Ten infant macaques (83.3%) successfully completed the gait protocol using the Catwalk apparatus at 28 days of life. The infants unable to obtain at least one usable run at day 21 and 28 were identified to be overall resistant to the task or distracted by the novel environment, which is consistent with the previous marmoset study (Pickett et al., 2020). As the infants aged, they became more interested in exploring the physical apparatus of the novel Catwalk environment impacting their performance on the task irrespective of their motor skills.

We identified that infants were already using a diagonal walking pattern at 14 days of life. This contrasts with previous literature that reported infant macaques use a lateral walking pattern at birth and transition into a diagonal walking pattern later on as they matured (Castell & Sackett, 1973; Dunbar, 1989; Hildebrand, 1967). Two animals did not conform to the definition of a diagonal walking pattern at day 28 of life. Both animals also had excluded runs due to being classified as running or jumping, which are more mature patterns. These infants may be walking faster or exploring more mature patterns of running/galloping contributing to the diagonal pattern not being utilized. At all timepoints, infants had variability in timing with about half of the infants using the diagonal pattern with a slight delay in timing where the front limb was landing a few seconds before the hind limb. Vilensky and colleagues (1983) had similar finding of the hind limbs being slightly delayed in landing time compared to the front limbs that was associated with how fast the infant was walking. This finding may show that early on infants are already utilizing a diagonal walking pattern but may have variable coordination as evidenced by variability in the timing when moving their contralateral limbs. The identification of this difference may be a novel result of using the Noldus Catwalk technology that allowed frame by frame analysis of the infants walking pattern as compared to previous observational assessment.

We did not find significant developmental changes from day 14 to day 28 in gait parameters, contrary to our original hypothesis. This study identified that there was only one significant change of phase dispersion LF→ LH in development between day 14 to day 28. We found a wide range of variability within and between animals at each timepoint for the individual Catwalk variables. This may further reflect the range of typical gait development for infant rhesus macaques. The data are consistent with human development where the first year of life is a variable, yet critical, time of motor development (Piek, 2002) and although there is an expected age to reach developmental milestones, many children reach them before or after while all falling within typical development (Zubler et al., 2022).

This study had several limitations. One limitation to this study was that all infant macaques were females. This was unplanned as the infants were enrolled in the study prior to knowing their sex. Research has found that males and females macaques have limited differences in the first month of life, but as they age males engage in more play behavior (Brown & Dixson, 2000). While the Noldus Catwalk is a promising tool for evaluating early gait in rhesus macaque, we did encounter challenges that require further evaluation and problem solving for future applications. One challenge that we encountered was the balance between number of trials and separating the infant from the mother. Ideally more trials collected over the first month of life would help to account for the high variability, but maternal separation creates stress, anxiety and increased vocalizations (Zhang et al., 2012). We limited separations to only occur at 14, 21, and 28 days of life to match with other neurodevelopmental testing and as recommended by animal care staff to support the mother-infant dyad attachment. Infant macaques may begin to walk as early as 3 days of life (Castell & Sackett, 1973) and may show early developmental differences. Future research may investigate gait development at 7 days of life, which might be a more sensitive timepoint for transitions between lateral to diagonal walking pattern. While the majority of rodent research has used task training to teach the rodent how to successfully walk across the Catwalk (Isvoranu et al., 2021), this was not attempted in this study due to only essential maternal separations occurring over the first month of life. The lack of training may have created difficulty for some animals due to the task novelty or possibly lack of understanding of task expectations.

Finally, the Catwalk apparatus and associated technology was designed for rodent models and there are limitations in the translation of this apparatus to larger animals. For instance, the length of the Catwalk limited the total number of step cycles a macaque could demonstrate in one run. By day 28 of life, most of the infants were only able to obtain the minimum number of footfalls required for analysis due to their size, whereas in the beginning of the study, they had additional step cycles that could be included in analyses. The size of the apparatus and embedded rodent gait parameters prevented standing on limb percentage from being automatically calculated and analyzed by the Noldus software. A longer walkway with a greater height would allow more footfalls to be captured to better understand rhesus macaque gait parameters at 28 days of life and potentially beyond the first month.

## 6. Conclusion

The Catwalk is a novel tool for early gait analysis that was successfully applied to provide detailed quantification of gait during the first month of life for rhesus macaques. Future applications should consider modifying the assessment based on challenges we encountered including adding additional timepoints, trying to reduce stress within the room, or modifying the length of the apparatus. Applying the Catwalk technology to infant rhesus macaque models studying neurodevelopmental deficits may provide a more detailed analysis of early gait differences and predict later developmental outcomes, through finding subtle differences in gait parameters.

Declaration of competing interest No conflict of interest

Data are available at: https://go.wisc.edu/4q8pb4.

## Acknowledgements

We would like to acknowledge the animal care, SPI, and veterinary staff for their support in caring for the infants included in this study, along with the staff and students of the Ausderau and Mohr labs for their continued contributions to and support of this work.

## Author credit statement

Sabrina A. Kabakov: Software, Validation, Formal analysis, Investigation, Data curation, Writing-original draft, Visualization, Project administration

Emma Crary: Data curation, Writing-original draft

Viktorie Menna: Investigation, Data curation, Project administration

Elaina Razo: Data curation

Jens Eickhoff: Formal analysis

Natalie Dulaney: Investigation, Data curation

Jack Drew: Data curation

Kathryn M. Bach: Investigation, Data curation, Project administration

Aubreonna M. Poole: Data curation

Madison Stumpf: Data curation

Ann M. Mitzey: Investigation

Kerri Malicki: Investigation, Resources, Project administration

Michele L. Schotzko: Investigation, Resources, Project administration

Kristen A. Pickett: Methodology

Nancy J. Schultz-Darken: Methodology

Marina E. Emborg: Methodology, Resources

David H. O’Connor: Funding acquisition

Thaddeus G. Golos: Funding acquisition

Emma L. Mohr: Conceptualization, Methodology, Supervision, Funding acquisition

Karla K. Ausderau: Conceptualization, Methodology, Validation, Investigation, Writing-original draft, Supervision, Project administration, Funding acquisition

All authors: Writing - review & editing

