## Supplemental Table 1 for "Quantification of Early Gait Development: Expanding the Application of Catwalk Technology to an Infant Rhesus Macaque Model"

Supplemental Materials:

**Supplemental Table 1.** Infant demographics

| Infant ID | Gestational age at inoculation (days) | Gestational age at birth (days) | Delivery method | Sex | Birth weight (kg) | Housing status | Days without a dam at 1 month | Age at timepoint 1 (days) | Age at timepoint 2 (days) | Age at timepoint 3 (days) |
|---|---|---|---|---|---|---|---|---|---|---|
| 044-507 | 48 | 163 | C-section | F | 0.5 | Surrogate | 6 | 13 | 20 | 27 |
| 044-505 | 42 | 161 | C-section | F | 0.434 | Surrogate | 4 | 15 | 22 | 29 |
| 044-506 | 48 | 160 | C-section | F | 0.563 | Nursery | 30 | 15 | 22 | 29 |
| 044-508 | 48 | 159 | Natural | F | 0.44 | Biological | 0 | 13 | 20 | 27 |
| 044-511 | 26 | 160 | C-section | F | 0.6 | Biological | 0 | 14 | 21 | 28 |
| 044-513 | 26 | 160 | C-section | F | 0.534 | Biological | 0 | 15 | 22 | 29 |
| 044-515 | 30 | 159 | C-section | F | 0.468 | Biological | 1 | 13 | 20 | 27 |
| 044-520 | 30 | 159 | C-section | F | 0.459 | Biological | 0 | 14 | 21 | 28 |
| 044-523 | 45 | 158 | C-section | F | 0.515 | Surrogate | 13 | 16 | 22 | 28 |
| 044-524 | 45 | 160 | C-section | F | 0.504 | Nursery | 30 | 16 | 22 | 29 |
| 044-525 | 45 | 160 | C-section | F | 0.484 | Nursery | 30 | 15 | 22 | 31 |
| 044-528 | 45 | 158 | C-section | F | 0.561 | Biological | 5 | 15 | 20 | 29 |
