## Supplemental Table 2 for "Quantification of Early Gait Development: Expanding the Application of Catwalk Technology to an Infant Rhesus Macaque Model"

**Supplemental Table 2.** Animal Catwalk Success

| Infant ID | Day 14 | Day 21 | Day 28 |
|---|---|---|---|
| 044-507 | 1 | 1 | 1 |
| 044-505 | 0 | 1* | 1 |
| 044-506 | 1 | 1 | 1 |
| 044-508 | 0 | 1 | 1* |
| 044-511 | 0 | 0 | 0 |
| 044-513 | 1 | 0 | 1 |
| 044-515 | 1 | 1 | 1 |
| 044-520 | 0 | 1 | 1 |
| 044-523 | 1 | 1 | 1 |
| 044-524 | 1 | 1 | 1 |
| 044-525 | 1* | 0 | 0 |
| 044-528 | 1 | 1 | 1 |
| Mean<br>(p-value#) | 66.7% | 75%<br>(.68) | 83.3%<br>(.42) |

1= at least 1 usable run was collected and 0= no usable runs. *= Equipment malfunction impacted background color preventing analysis of the run. #= p-value comparison to day 14.
