## Supplemental Table 3 for "Quantification of Early Gait Development: Expanding the Application of Catwalk Technology to an Infant Rhesus Macaque Model"

**Supplemental Table 3.** Test-retest variability. Intra-class correlation Coefficient (ICC)

| Gait Variable | Day 14 | Day 21 | Day 28 |
|---|---|---|---|
| Speed (cm/s) | 0.19 | **0.50** | 0.20 |
| RF Stand Index | -0.17 | **0.56** | -0.27 |
| RH Stand Index | 0.18 | 0.08 | 0.39 |
| LF Stand Index | 0.08 | 0.19 | -0.03 |
| LH Stand Index | 0.03 | **0.63** | 0.24 |
| RF Duty Cycle (%) | -0.09 | **0.51** | 0.26 |
| RH Duty Cycle (%) | 0.34 | 0.00 | -0.24 |
| LF Duty Cycle (%) | -0.01 | 0.46 | 0.06 |
| LH Duty Cycle (%) | 0.16 | -0.03 | 0.40 |
| RF Swing (%) | -0.08 | **0.51** | 0.26 |
| RH Swing (%) | 0.37 | 0.02 | -0.25 |
| LF Swing (%) | -0.04 | 0.47 | 0.06 |
| LH Swing (%) | 0.12 | -0.03 | 0.37 |
| RF Single Stance (%) | -0.09 | 0.35 | 0.05 |
| RH Single Stance (%) | -0.04 | -0.02 | 0.01 |
| LF Single Stance (%) | -0.09 | **0.50** | 0.26 |
| LH Single Stance (%) | **0.54** | 0.33 | -0.23 |
| Front Dual Stance (%) | 0.29 | **0.63** | 0.24 |
| Hind Dual Stance (%) | 0.49 | **0.55** | 0.00 |
| RF Stride Length (cm) | 0.45 | **0.55** | 0.21 |
| RH Stride Length (cm) | **0.57** | **0.60** | 0.13 |
| LF Stride Length (cm) | **0.53** | 0.48 | 0.39 |
| LH Stride Length (cm) | **0.58** | **0.54** | 0.13 |
| Front Base of Support (cm) | 0.42 | 0.29 | 0.35 |
| Hind Base of Support (cm) | 0.48 | 0.03 | 0.33 |
| Right Print Position (cm) | -0.21 | 0.00 | -0.13 |
| Left Print Position (cm) | -0.18 | 0.49 | -0.02 |
| Standing on Diagonal Limbs (%) | **0.66** | -0.16 | -0.16 |
| Standing on Three Limbs (%) | **0.52** | 0.01 | -0.13 |
| Standing on Four Limbs (%) | -0.01 | **0.66** | 0.36 |
| Average of All | 0.20 | 0.32 | 0.11 |

ICC <.5 = poor reliability, .5-.75= moderate reliability, >.75= good/excellent reliability. Bold indicates ICC with moderate reliability.
