## Supplemental Table 4 for "Quantification of Early Gait Development: Expanding the Application of Catwalk Technology to an Infant Rhesus Macaque Model"

**Supplemental Table 4.** Catwalk parameters comparison between day 14, 21, and 28 days of life

| Gait variable | Limbs | Day 14-Day 21 | Day 14- Day 28 | Day 21- Day 28 |
|---|---|---|---|---|
| | | *Speed* | | |
| Speed (cm/s) | | **0.04989** | 0.87461 | **0.03296** |
| Stand Index | RF | 0.11727 | 0.8267 | 0.16851 |
| | LF | 0.73413 | 0.78848 | 0.50342 |
| | RH | 0.19356 | 0.94367 | 0.21992 |
| | LH | 0.06931 | 0.77733 | 0.11674 |
| | | *Dynamic paw measurements* | | |
| Duty Cycle (%) | RF | 0.36731 | 0.938 | 0.31496 |
| | LF | 0.14278 | 0.77254 | 0.07602 |
| | RH | 0.07793 | 0.90677 | 0.09712 |
| | LH | 0.19514 | 0.24012 | 0.90896 |
| Swing (%) | RF | 0.48864 | 0.87345 | 0.38233 |
| | LF | 0.2336 | 0.57706 | 0.07834 |
| | RH | 0.10318 | 0.87968 | 0.13428 |
| | LH | 0.18908 | 0.29696 | 0.90901 |
| Single Stance (%) | RF | 0.45251 | 0.30446 | 0.0899 |
| | LF | 0.43515 | 0.98969 | 0.4172 |
| | RH | 0.32459 | 0.24634 | 0.75181 |
| | LH | 0.17223 | 0.95977 | 0.14837 |
| Dual Stance (%) | Front | 0.21547 | 0.41249 | **0.03479** |
| | Hind | 0.07987 | 0.46249 | 0.35989 |
| | | *Static paw measurements* | | |
| Stride Length (cm) | RF | **0.04914** | 0.34242 | 0.35813 |
| | LF | **0.01882** | 0.15325 | 0.44256 |
| | RH | 0.14301 | 0.83412 | 0.21041 |
| | LH | 0.20951 | 0.51203 | 0.65976 |
| Base of | Front | 0.91034 | 0.98089 | 0.88549 |

| | | | | |
|---|---|---|---|---|
| Support (cm) | Hind | 0.08815 | 0.27714 | 0.70118 |
| Print Position (%) | Right | 0.12141 | 0.2084 | 0.80175 |
| | Left | 0.54636 | 0.52074 | 0.85922 |
| *Inter-paw coordination* | | | | |
| Phase Dispersion (%) | RF->LH | 0.78031 | 0.33418 | 0.43982 |
| | LF->RH | 0.25166 | 0.06377 | 0.27224 |
| | LH->RH | 0.77638 | 0.99032 | 0.75448 |
| | LF->RF | 0.8772 | 0.44307 | 0.3395 |
| | RF->RH | 0.67086 | 0.47286 | 0.256 |
| | LF->LH | 0.4667 | **0.0002** | 0.05667 |
| Standing on (%) | diagonal | 0.9216 | 0.83714 | 0.74023 |
| | three | 0.6842 | 0.87043 | 0.81626 |
| | four | 0.55379 | 0.99875 | 0.54597 |
| Walking pattern (%) | Diagonal | 0.97258 | 0.2125 | 0.3998 |

Bold indicates p-values less than 0.05.
